# The role of mitochondrial complex I in the proinflammatory response to polylactide implants

**DOI:** 10.1101/2024.08.12.607680

**Authors:** Chima V. Maduka, Ashley V. Makela, Anthony Tundo, Evran Ural, Katlin B. Stivers, Mohammed Alhaj, Ramani Narayan, Stuart B. Goodman, Nureddin Ashammakhi, Jennifer H. Elisseeff, Kurt D. Hankenson, Christopher H. Contag

## Abstract

During the foreign body response, immune cells are metabolically rewired after exposure to breakdown products of various biomaterials, including polylactide (PLA) and polyethylene. Particles of polyethylene interact with Toll-like receptor 4 on macrophages, resulting in increased oxygen consumption that forms reactive oxygen species at complex I of the mitochondrial electron transport chain (mETC). However, PLA degradation products bind to monocarboxylate transporters for downstream signaling with elevated oxygen consumption rates, whose functional implication is unclear and remains inferred from cellular responses to polyethylene biomaterials. By chemically probing the function of the mETC, we show that proinflammatory macrophages activated by exposure to amorphous PLA (aPLA) breakdown products rely on mitochondrial respiration for ATP production, independent of oxygen consumption rates. In contrast, macrophages activated by semi-crystalline PLA (cPLA) breakdown products exhibit a metabolic phenotype wherein ATP levels are unaffected by changing oxygen consumption rates. In subcutaneous implants, the incorporation of metformin in aPLA or cPLA to chemically inhibit complex I did not effectively modulate the proinflammatory response to biomaterials, suggesting that PLA degradation products elicit a distinct metabolic program, thus providing an alternative perspective on the role of mitochondrial respiration in the inflammatory response to biomaterials.

## Introduction

During the foreign body response to polylactide (PLA), activated immune cells exhibit altered metabolic profiles, including simultaneously increased glycolytic flux and oxygen consumption rates^1^. These metabolic phenotypes are accompanied by elevated adenosine triphosphate (ATP) levels and affected by the chirality of PLA implants^2^. In human patients, aberrant glycolytic flux around PLA implants manifests as higher F-18 fluorodeoxyglucose uptake during positron emission tomography scans^3,4^, an observation reproduced in rodent models^1^. Prevailing immunometabolic signals around biomaterials affect the trafficking of monocytes from the circulation into the microenvironment surrounding the biomaterial in a process dependent on CCR2 and CX3CR1 signaling^5^. Elevated glycolytic flux and altered bioenergetics (ATP levels) induced by PLA degradation products result in increased production of proinflammatory cytokines and chemokines, including IL-6, TNF-α, IL-1β and CCL2; accordingly, targeting different steps in the glycolytic pathway modulates bioenergetics and proinflammatory protein levels^1^. In-vivo, control of glycolysis by embedding inhibitors in implants reduces the numbers of recruited neutrophils and regulates the relative composition and activation states of monocytes, macrophages and dendritic cells^5^. While the ratios of proinflammatory (CD86^+^CD206^-^) to anti-inflammatory (CD206^+^) cells are reduced, the ratios of anti-inflammatory and transition (CD86^+^CD206^+^) cells to proinflammatory populations are increased, creating a pro-regenerative environment where IL-4 levels from γd+ T cells and T helper 2 cells may increase^5^. As with PLA degradation products, glycolytic reprogramming drives chronic inflammation from sterile and endotoxin-contaminated polyethylene wear particles as well as hydroxyapatite particles and composites, playing a pivotal role in the longevity of total joint arthroplasties^6-10^.

While the functional implication of altered glycolysis in the PLA microenvironment is understood, it is unclear what role mitochondrial respiration (oxidative phosphorylation) plays in the proinflammatory response to implanted PLA. Mitochondrial respiration has been exploited using biomaterials to induce cellular biostasis^11^. Furthermore, increased oxygen consumption at complex I of the mitochondrial electron transport chain (mETC) has been shown to result in elevated superoxide production in immune cells, following ischemia-reperfusion injury^12^ and in cancer biology, where chronic inflammation has become a hallmark of cancer progression^13^. Through reverse electron transport, mitochondrial membrane potential sustains the generation of mitochondrial reactive oxygen species (ROS)^14,15^, a phenomenon observed in various types of cancers^16^, neuroinflammation^17-19^ and inflammation induced by endotoxin^20^ or polyethylene particles^21^. Consequently, the pharmacologic inhibition of oxygen consumption at complex I by rotenone or metformin^21,22^ suppresses proinflammatory activation by endotoxin or polyethylene particles. Immunomodulatory effects are likely the result of increased oxygen consumption enabling neutrophil activation by increasing ROS levels^23^; underlying the ability of macrophage and dendritic cells to activate T cells by antigen presentation^24^; and sustaining antigen-specific expansion of T cells, independent of glycolytic machinery^25^. Based on these observations, we hypothesized that macrophages exposed to PLA breakdown products manifest increased oxygen consumption rates for inflammatory activation, and this process is regulated by the mitochondrial electron transport chain (mETC); therefore, modulating the mETC will reduce inflammation in the PLA microenvironment.

Contrary to this hypothesis, our current data suggest that proinflammatory macrophages activated by exposure to amorphous PLA (aPLA) breakdown products rely on mitochondrial respiration for ATP production, independent of oxygen consumption rates. In contrast, crystalline PLA (cPLA) breakdown products elicit a metabolic phenotype wherein ATP levels are unaffected by changing oxygen consumption rates in activated macrophages. Inhibition of oxygen consumption at complex I of the mETC was accomplished by rotenone and metformin after exposure of macrophages to cPLA but not aPLA breakdown products. Overall, the incorporation of metformin into aPLA or cPLA did not effectively modulate the proinflammatory response to implants. Proinflammatory macrophage and dendritic cell proportions were not decreased nor were anti-inflammatory populations increased in aPLA or cPLA implants containing metformin. However, the incorporation of metformin into cPLA implants increased Arginase 1 levels among macrophages, dendritic cells and class II major histocompatibility complex (MHC II) expressing dendritic cells. Around these cPLA implants, there were concomitantly elevated IFN-γ and IL-4 levels from T helper cells. These results contrast the strong immunomodulatory effects observed by metformin’s inhibition of complex I when immune cell activation occurs subsequent to Toll-like receptor 4 signaling, suggesting that PLA degradation products elicit a distinct metabolic program during proinflammatory activation that may be affected by the crystallinity of the biomaterial. Elucidating the underpinnings of metabolic rewiring around biodegradable biomaterials will inform the rational design of implants applied in regenerative medicine^26-29^.

## Results

Using a bioenergetic model that we had previously developed and validated^1,2^, we exposed primary bone marrow-derived macrophages (BMDMs) to amorphous PLA (aPLA) or semi-crystalline PLA (cPLA) breakdown products (also called extracts), having validated the physicochemical properties of aPLA and cPLA^1^. To chemically probe the function of complex I, III and V of the mETC, we treated macrophage conditions with their respective inhibitors, including rotenone, metformin, antimycin A and oligomycin. Rotenone inhibits complex I of the mETC, an effect reproduced by metformin, which is a Food and Drug Administration (FDA)-approved drug prescribed for diabetic patients^21,22^. Antimycin A and oligomycin inhibit complex III and V, respectively, of the mETC^21^. Compared to untreated controls, exposure to aPLA or cPLA breakdown products increased ATP levels (Fig. 1a-b)^1^. While the inhibition of complex I, III and V decreased ATP levels in the aPLA-treated groups, there were no significant reductions with cPLA groups (Fig. 1a-b). Interestingly, with both rotenone and metformin, inhibition of complex I was dose-dependent in the aPLA-treated group (Fig. 1a). Inhibition of complex I similarly modulates ATP production in other inflammatory conditions, such as pancreatic cancer, where signaling through the HMGB1/RAGE inflammatory pathway increases ATP levels^30^. Observed bioenergetic changes were not associated with significant changes in cell viability, suggesting minimal toxicity (Fig. 2a-b). In contrast, glycolytic inhibition of cells exposed to aPLA or cPLA reduces ATP levels and is accompanied by mild decreases in cell numbers^1^.

**Figure 1.**
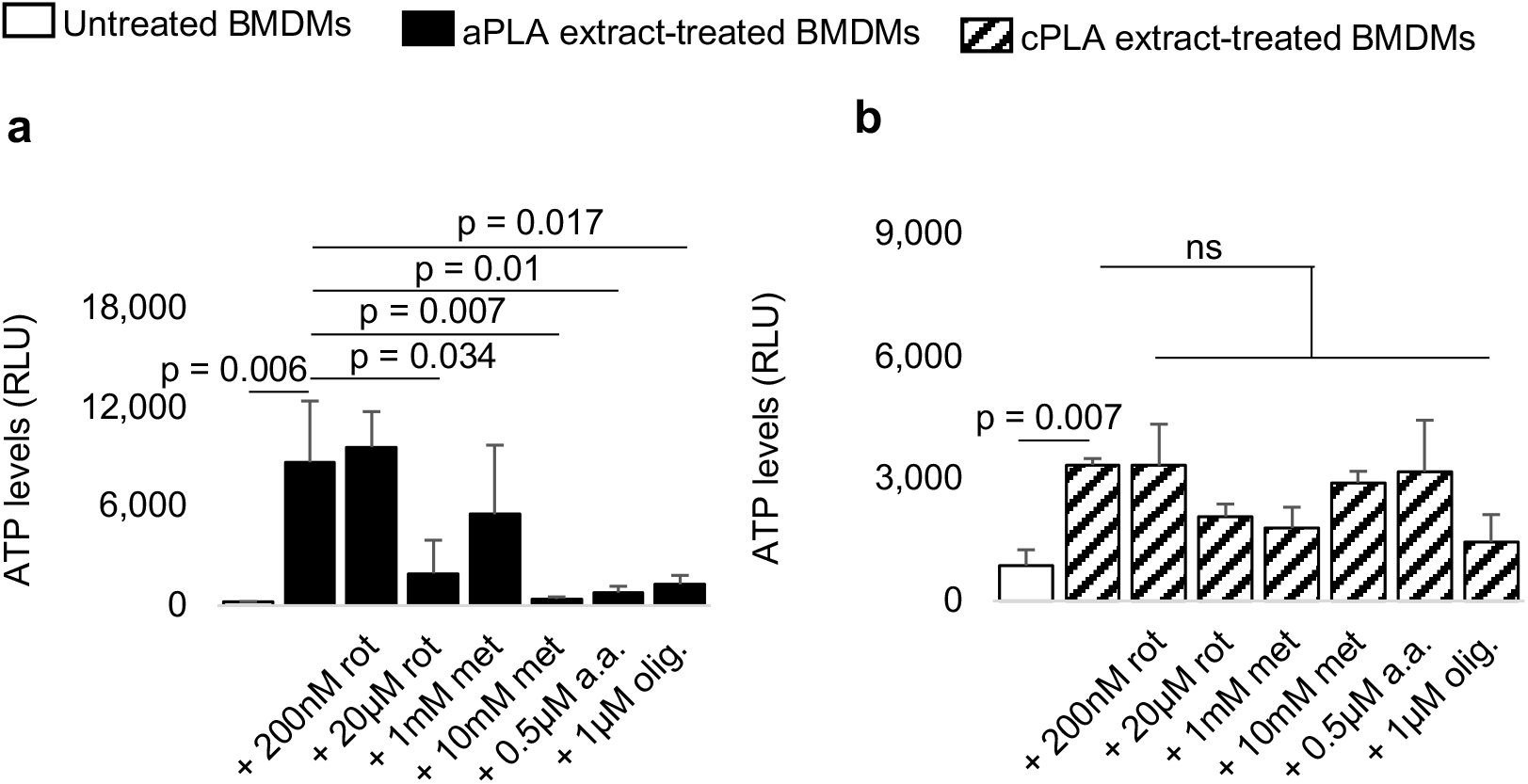
Inhibition of mitochondrial respiration differentially affects bioenergetics (ATP levels) in primary bone marrow-derived macrophages (BMDMs) exposed to polylactide (PLA) degradation products (extracts). **a**, Compared to untreated BMDMs, amorphous PLA (aPLA) extracts increase ATP levels; elevated bioenergetics is reduced by inhibiting of the mitochondrial electron transport chain using rotenone (rot), metformin (met), antimycin A (a.a.) and oligomycin (olig.). **b**, Increased bioenergetics from exposure to crystalline PLA (cPLA) extracts is not decreased by inhibition of mitochondrial respiration. Not significant (ns), mean (SD), n = 3, one-way ANOVA followed by Tukey’s post-hoc test; 100 μl aPLA or 150 μl cPLA extract with corresponding controls were used after 7 days in-culture.

**Figure 2.**
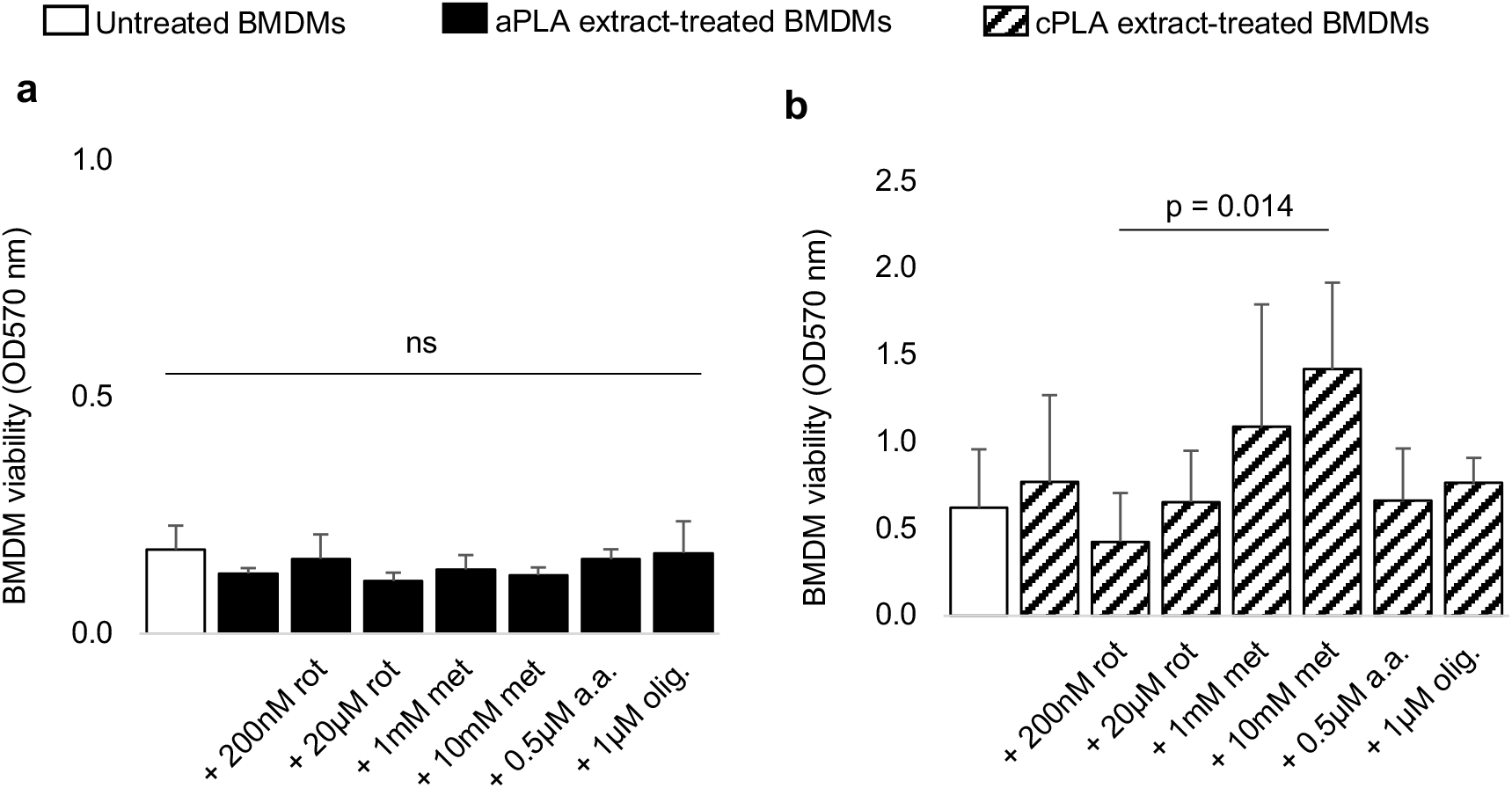
Inhibition of the mitochondrial electron transport chain does not reduce cell viability. **a-b**, Compared to untreated primary bone marrow-derived macrophages (BMDMs), exposure to amorphous polylactide (aPLA; **a**) or crystalline polylactide (cPLA; **b**) does not affect cell numbers; treatment with inhibitors of mitochondrial respiration, including rotenone (rot), metformin (met), antimycin A (a.a.) and oligomycin (olig.) does not reduce cell viability relative to PLA-treated cells. Not significant (ns), mean (SD), n = 5, one-way ANOVA followed by Tukey’s post-hoc test; 100 μl aPLA or 150 μl cPLA extract with corresponding controls were used after 7 days in-culture.

There was increased oxygen consumption rate (OCR) in BMDMs exposed to cPLA breakdown products, which was decreased by chemically inhibiting complex I or V, but not complex III (Fig. 3a). Elevated OCR related to the mETC drives inflammatory activation to polyethylene particles^21^, which play a pivotal role in chronic inflammation underlying implant failure after total knee and hip replacements^6-10^. Among cancers, where chronic inflammation is a hallmark of cancer progression^13^, elevated OCR has been noted in oral squamous cell carcinoma^31^, triple-negative breast cancer^32^ and pancreatic cancer^30^. With pancreatic cancer, ATP levels are reduced by chemical inhibition of complex I via rotenone’s ability to modulate oxygen consumption at complex I of the mETC^30^. Interestingly, inhibition of the mETC did not decrease OCR in untreated BMDMs and BMDMs exposed to aPLA breakdown products (Fig. 3b-c). Together with the data on ATP levels, these results suggest that ATP production may be uncoupled from oxygen consumption through the mETC, distinct from ATP generation in resting cells under physiological conditions^33^. That changes in OCR were observed with crystalline (Fig. 3a) and not amorphous (Fig. 3b) PLA degradation products suggest a role for crystallinity, in addition to stereochemistry^2^, as a factor affecting metabolic changes in activated macrophages exposed to biomaterials. Previously, in a dose-dependent manner, glycolytic inhibition consistently reduced OCR in macrophages exposed to various polylactide breakdown products^1,2^, suggesting that glycolysis may be coupled to the mETC in macrophages activated by polylactide breakdown products.

**Figure 3.**
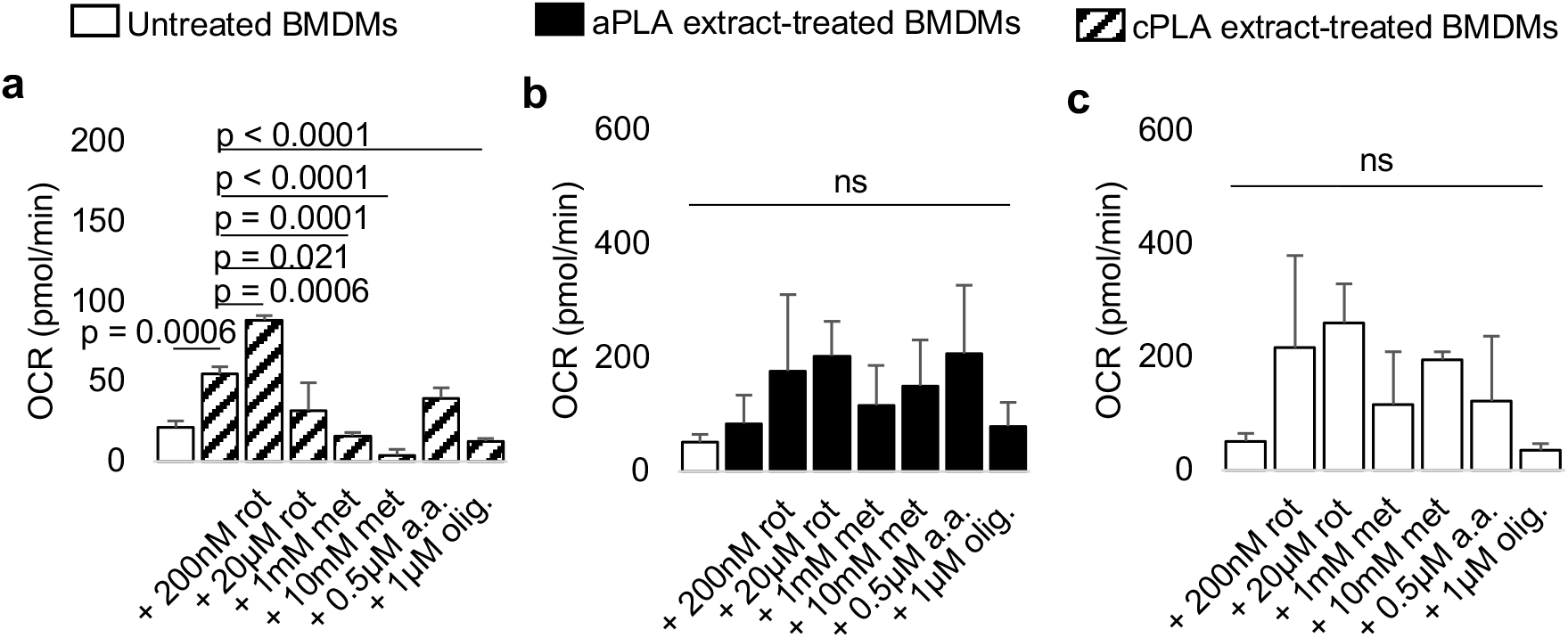
Pharmacologically targeting complex I of the electron transport chain (ETC) reduces oxygen consumption rate (OCR) in primary bone marrow-derived macrophages (BMDMs) exposed to crystalline but not amorphous polylactide (PLA) degradation products (extracts). **a-b**, In BMDMs exposed to crystalline PLA (cPLA; **a**) but not amorphous PLA (aPLA; **b**) extracts, elevated OCR is decreased by inhibition of complex I using rotenone (rot) or metformin (met), or complex V using oligomycin (olig.), but not complex III using antimycin A (a.a.). **c**, Targeting the ETC using rot, met, a.a. and olig. in untreated BMDMs does not decrease OCR. Mean (SD), n = 3 in, one-way ANOVA followed by Tukey’s post-hoc test; 100 μl aPLA or 150 μl cPLA extract with corresponding controls were used after 7 days in-culture.

Although inhibition of complex I was selectively dose-dependent with cPLA but not aPLA extracts (Fig. 3a-b), we sought to determine whether there was a relationship between oxygen consumption at complex I and mitochondrial reactive oxygen species (ROS). Oxygen consumption at complex I has been shown to be a driver of mitochondrial ROS with sterile polyethylene particles^21^ as well as bacterial endotoxin^20^, where mitochondrial membrane potential underpins mitochondrial ROS production through reverse electron transport^20^. Exposed macrophages were stained with MitoSOX red (for mitochondrial ROS) gated on MitoTracker Green (for mitochondria). Surprisingly, with respect to the proportion of macrophages producing mitochondrial ROS, there was no difference between untreated controls and macrophage exposed to aPLA extracts (Fig. 4a-b). Interestingly, cPLA extracts reduced the proportion of macrophages producing mitochondrial ROS, and the addition of metformin to aPLA and cPLA extracts produced opposite results (Fig. 4c-e), corroborated by quantitative dataset (Fig. 4f). Together this data suggest that oxygen consumption in macrophages exposed to aPLA and cPLA extracts may be unrelated to mitochondrial ROS production, distinct from findings involving polyethylene particles^21^, endotoxin^20^ and neuroinflammation^17-19^. Despite these data, metformin reduced levels of proinflammatory cytokines and chemokines, including IL-1β, IL-6, MCP-1 and TNF-α (with TNF-α, reduction occurred with only aPLA extract), in macrophages activated by aPLA and cPLA extracts (Fig. 5a-d). Levels of anti-inflammatory cytokines, such as IL-10 and IL-4, were either reduced or unaffected by the addition of metformin in cultured macrophages (Fig. 5e-f), and we could not detect IFN-γ and 1L-13 levels (data not shown). While similar in its effects to glycolytic inhibition, unlike metformin’s effects on anti-inflammatory cytokines, glycolytic inhibition tend to increase IL-10 expression among macrophages activated by polylactide extracts^1^.

**Figure 4.**
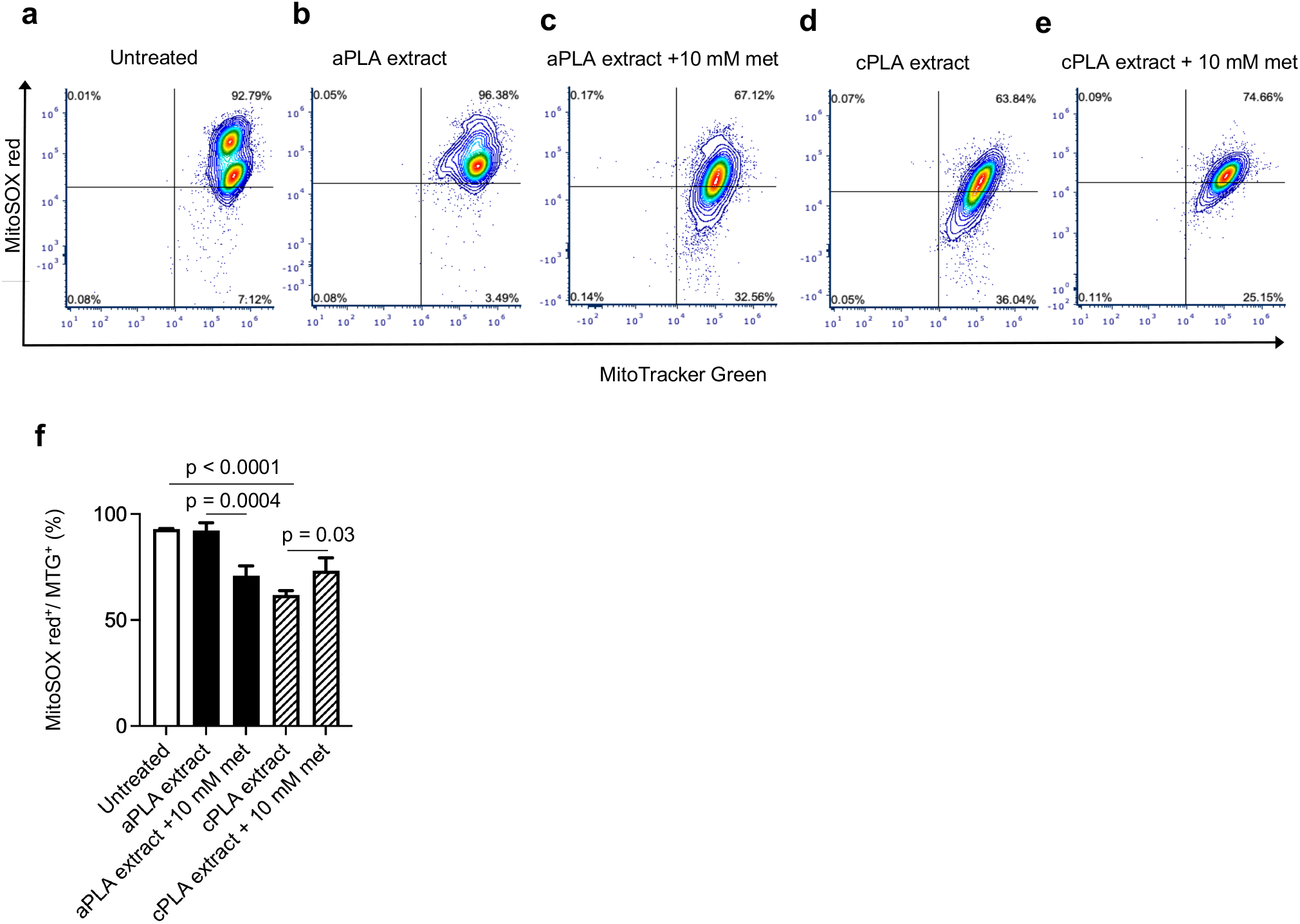
Differential mitochondrial reactive oxygen species (ROS) production occurs with polylactide (PLA) degradation products. **a-c**, Compared to untreated controls (**a**), representative flow plots suggest that macrophages exposed to amorphous PLA (aPLA; **b**) extracts exhibit unaltered levels of MitoSOX red (mitochondrial ROS), with metformin (met; **c**) decreasing mitochondrial ROS levels. **d-e**, In comparison to untreated controls, prolonged exposure of macrophages to crystalline PLA (cPLA; **d**) extracts reduce mitochondrial ROS, but the addition of metformin increases mitochondrial ROS levels (**e**). **f**, Quantified proportion of mitochondria (MitoTracker Green, MTG) that produce mitochondrial ROS corroborate flow cytometry plots. Mean (SD), n = 3, one-way ANOVA followed by Tukey’s post-hoc test; 100 μl aPLA or 150 μl cPLA extract with corresponding controls were used after 7 days in-culture.

**Figure 5.**
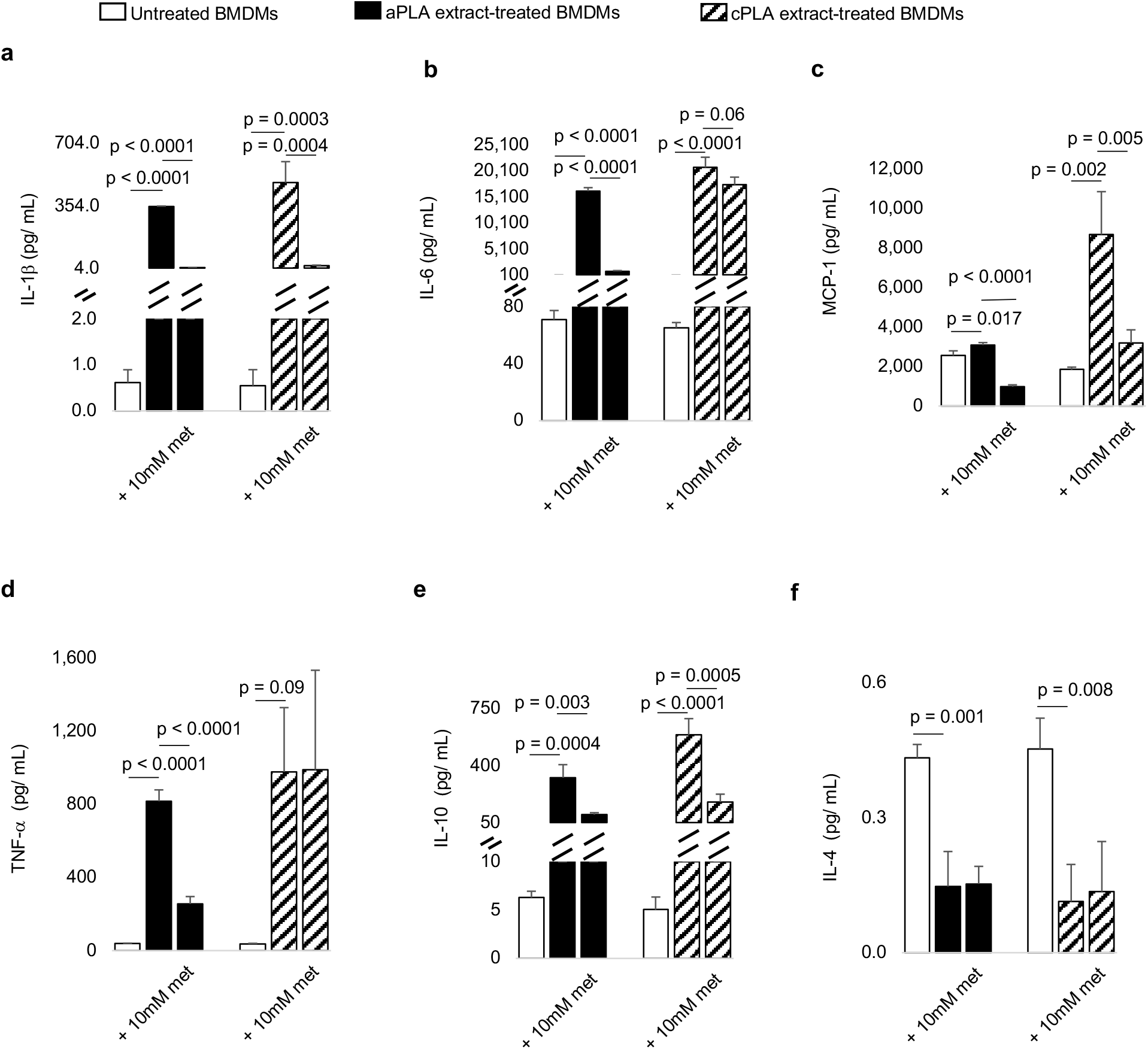
Inhibition of mitochondrial respiration reduces proinflammatory activation of primary bone marrow-derived macrophages exposed to polylactide (PLA) degradation products (extracts). **a-c**, In BMDMs, proinflammatory cytokine (protein) levels, including IL-1β (**a**), IL-6 (**b**) and MCP-1 (**c**) are increased by both amorphous PLA (aPLA) and crystalline PLA (cPLA) extracts; inhibition of the mitochondrial electron transport chain by metformin (met) reduces activation of BMDMs. **d**, In BMDMs exposed to aPLA, increased TNF-α protein expression is decreased by metformin. **e-f**, Whereas aPLA and cPLA extracts increase IL-10 protein expression (**e**), they decrease IL-4 levels (**f**); addition of metformin does not increase either anti-inflammatory cytokine relative to PLA alone. Mean (SD), n = 3 in all except the cPLA group for TNF-α where n = 2 as one value which exceeded the threshold was excluded, one-way ANOVA followed by Tukey’s post-hoc test; 100 μl aPLA or 150 μl cPLA extract with corresponding controls were used after 7 days in-culture.

To test for potential immunomodulatory effects arising from the inhibition of complex I of the mETC, we used a subcutaneous implantation model to compare reprocessed aPLA and cPLA to respective polymers that incorporated metformin by melt-blending. In comparison to sham groups, nucleated hematopoietic (CD45^+^) and macrophage (F4/80^+^) cell populations were increased around aPLA and cPLA biomaterials (Fig. 6a-b)^5^. However, the incorporation metformin did not reduce CD45^+^ or F4/80^+^ populations (Fig. 6a-b). We assigned proinflammatory and anti-inflammatory populations as CD86^+^CD206^-^ and CD206^+^, respectively^34,35^. Whereas the fold change of proinflammatory with respect to anti-inflammatory macrophages was increased, the fold change of anti-inflammatory with respect to proinflammatory macrophages was decreased in the aPLA and cPLA microenvironment (Fig. 6c-d)^5^. The incorporation of metformin neither decreased proinflammatory nor increased anti-inflammatory ratios (Fig. 6c-d). This is in contrast with glycolytic inhibition, where proinflammatory ratios are decreased concomitant with an increase in anti-inflammatory levels in aPLA implants and composites^5,10^. Compared to sham groups, arginase 1 (Arg1) levels were increased in the biomaterial microenvironment of aPLA and cPLA implants (Fig. 6e). Interestingly, the incorporation of metformin increased Arg1 levels in the cPLA but not the aPLA microenvironment (Fig. 6e-j).

**Figure 6.**
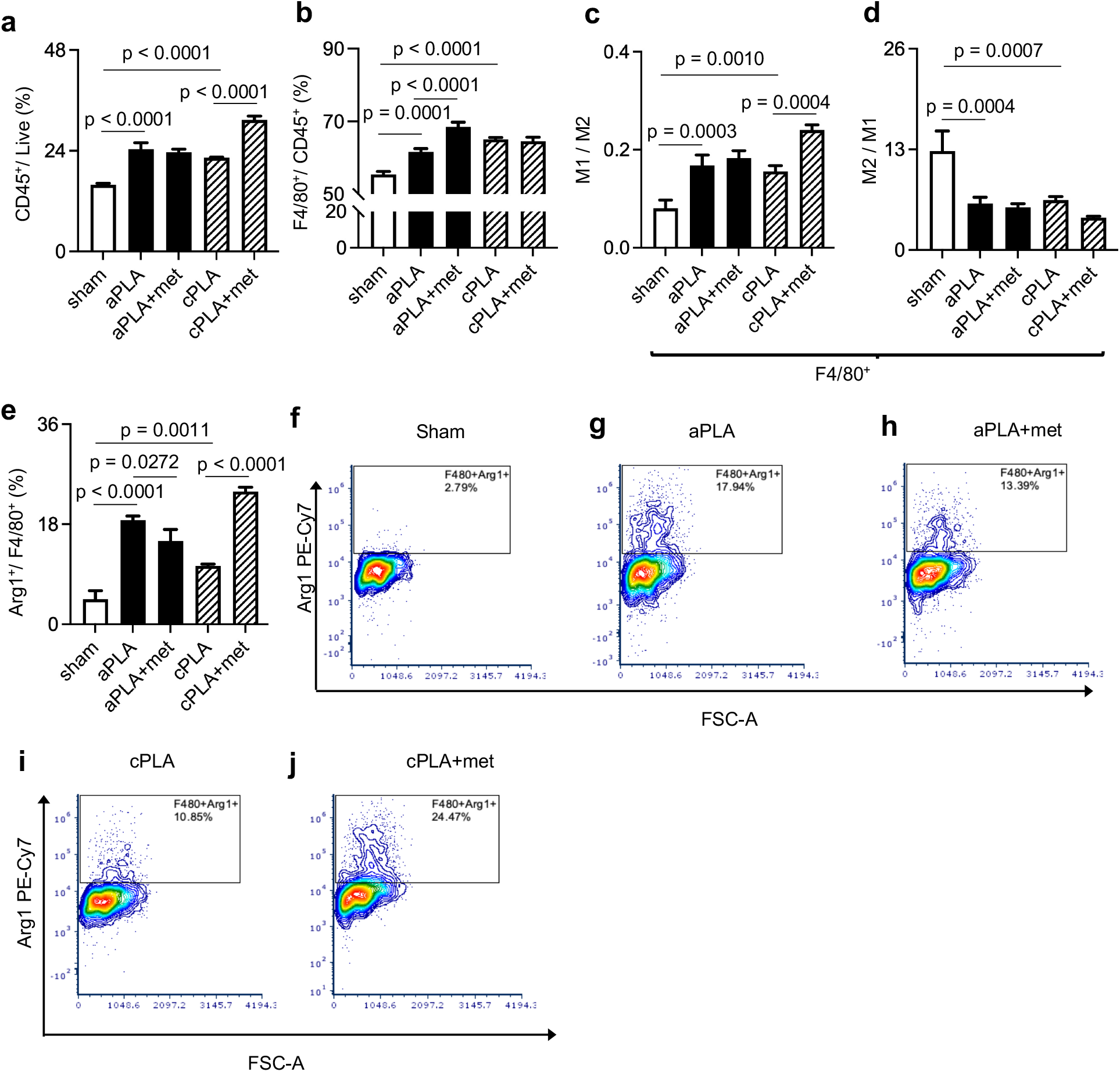
The frequency and proinflammatory states of macrophages are not reduced by incorporation of metformin in polylactide (PLA) implants. **a**, Unlike with amorphous PLA (aPLA), the incorporation of metformin (met) in crystalline PLA (cPLA) increases the frequency of nucleated hematopoietic (CD45^+^) populations in the implant microenvironment. **b**, With the incorporation of metformin, there is increased macrophage recruitment in the aPLA and not the cPLA microenvironment. **c**, The fold change of proinflammatory (M1; CD86^+^CD206^-^) with respect to anti-inflammatory (M2; CD206^+^) macrophage is neither reduced in the aPLA nor cPLA microenvironment following the incorporation of metformin. **d**, The fold change of M2 to M1 macrophage is unchanged by the incorporation of metformin. **e-j**, Unlike with aPLA, arginase 1 (Arg1) levels are increased by the incorporation of metformin in the cPLA microenvironment (e) as shown in representative flow cytometry plots (f-j). Mean (SD), one-way ANOVA followed by Tukey’s multiple comparison test, n = 3.

Dendritic (CD11c^+^) cell populations were elevated in the cPLA but not the aPLA microenvironment (Fig. 7a)^5^. The incorporation of metformin in either aPLA or cPLA increased dendritic cell numbers compared to aPLA or cPLA alone (Fig. 7a). The fold change of proinflammatory with respect to anti-inflammatory dendritic cells was increased around aPLA and cPLA biomaterials (Fig. 7b)^5^. Remarkably, the incorporation of metformin only modestly decreased proinflammatory ratios (Fig. 7b-g), unlike the profound effects observed with glycolytic inhibitors^5,10^. We observed that the fold change of anti-inflammatory with respect to proinflammatory dendritic cells was decreased around aPLA and cPLA biomaterials (Fig. 7h)^5^. In contrast to the effects of glycolytic inhibition^5,10^, the incorporation of metformin in aPLA or cPLA implants had no effects on anti-inflammatory ratios (Fig. 7h). Consistent with observations made with macrophages, the incorporation of metformin in cPLA increased Arg1 levels compared to cPLA alone (Fig. 7i).

**Figure 7.**
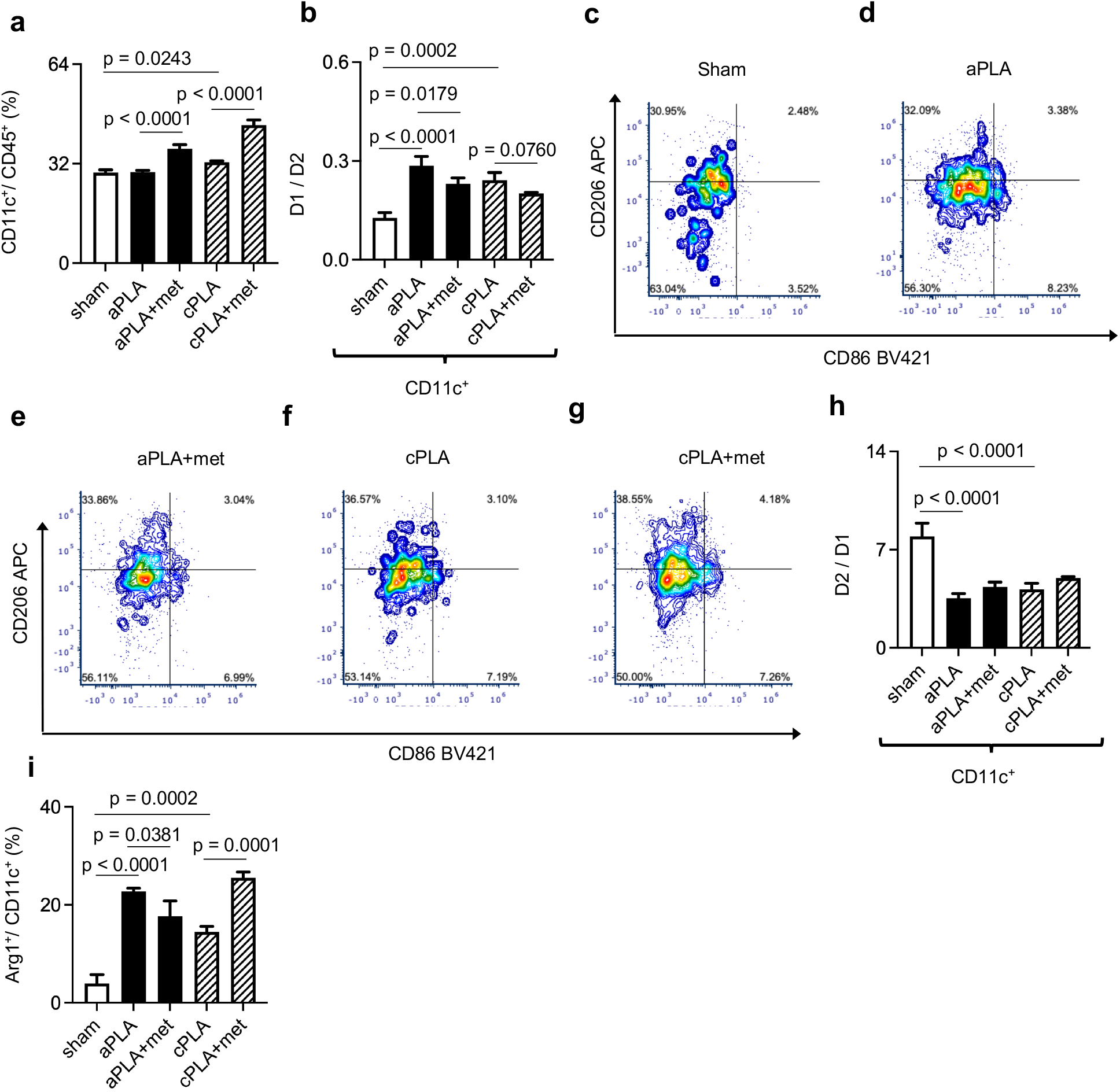
The incorporation of metformin increases the proportion and affects the polarization states of dendritic cells in the polylactide (PLA) microenvironment. **a**, The incorporation of metformin increases the proportion of recruited dendritic (CD11c+) cells in the amorphous PLA (aPLA) and crystalline PLA (cPLA) implant microenvironment. **b-g**, The fold change of proinflammatory (D1; CD86^+^CD206^-^) dendritic cells with respect to anti-inflammatory (D2; CD206^+^) dendritic cells is deceased with incorporation of metformin in the aPLA microenvironment (b) as shown in representative flow plots (c-g); this difference is not significant with cPLA (b-g). **h**, The fold change of D2 to D1 dendritic cells is unchanged by the incorporation of metformin in aPLA or cPLA implants. **i**, Unlike with aPLA, Arginase 1 (Arg1^+^) dendritic cell populations are increased by the incorporation of metformin in cPLA implants. Mean (SD), one-way ANOVA followed by Tukey’s or Newman-Keul’s multiple comparison test, n = 3.

We observed that dendritic cells expressing class II major histocompatibility complex (MHC II) molecules were decreased in the aPLA but not the cPLA microenvironment when compared to sham controls (Fig. 8a)^5^. With either aPLA or cPLA, the incorporation of metformin increased the frequency of dendritic cells expressing MHCII when compared to aPLA or cPLA alone (Fig. 8a), an observation also made with the incorporation of glycolytic inhibitors^5,10^. The fold change of proinflammatory with respect to anti-inflammatory dendritic cells expressing MHCII was increased around aPLA and cPLA biomaterials (Fig. 8b)^5^. Additionally, the incorporation of metformin reduced proinflammatory levels of dendritic cells expressing MHCII in the aPLA but not the cPLA groups (Fig. 8b). In contrast, the incorporation of metformin in aPLA or cPLA implants had no effects on the fold change of anti-inflammatory with respect to proinflammatory dendritic cells expressing MHCII (Fig. 8c). The incorporation of metformin in cPLA increased Arg1 levels among dendritic cells expressing MHCII compared to cPLA alone (Fig. 8d). We observed that CD3^+^ T cell populations were decreased around aPLA and cPLA implants, and that the inclusion of metformin did not alter T cell frequencies (Fig. 8e). Among T cells, CD8^+^ cytotoxic T cells as well as CD4^+^ T helper cells were increased around aPLA and cPLA implants, but the addition of metformin did not alter CD8 and CD4 levels (Fig. 8f-g). Interestingly, the inclusion of metformin in cPLA but not aPLA implants concomitantly elevated IL-4 and IFN-γ cytokine production from T helper cells (Fig. 8h-i).

**Figure 8.**
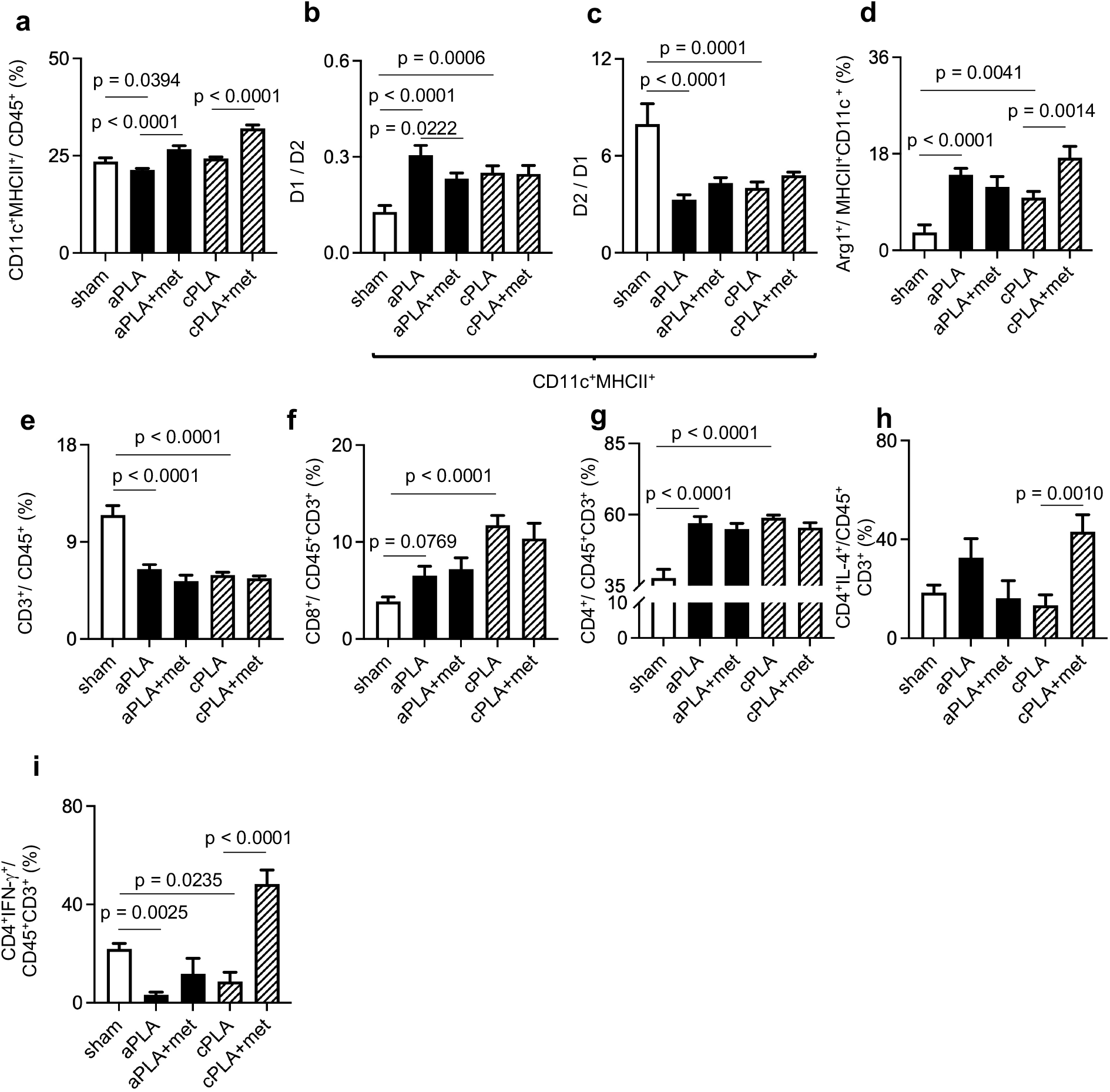
Metformin exerts differential effects on the activation states of dendritic cells expressing class II major histocompatibility complex (MHC II) molecules and T-cells in the polylactide (PLA) microenvironment. **a**, The frequency of dendritic cells expressing MHC II is increased by the incorporation of metformin (met) in amorphous PLA (aPLA) and crystalline PLA (cPLA) implants. **b**, Unlike with cPLA implants, the incorporation of metformin in aPLA implants reduces the fold change of proinflammatory (D1; CD86^+^CD206^-^) with respect to anti-inflammatory (D2; CD206^+^) dendritic cells expressing MHC II. **c**, The fold change of D2 with respect to D1 dendritic cells expressing MHC II is unchanged by the incorporation of metformin in aPLA or cPLA implants. **d**, Unlike with aPLA implants, the incorporation of metformin in cPLA implants increases the frequency of Arginase 1 (Arg1^+^) dendritic cells expressing MHC II. **e**, Overall T-cell (CD3^+^CD11b^-^ gated on CD45^+^ cells) populations are unchanged by the incorporation of metformin in aPLA or cPLA implants. **f-g**, Frequencies of cytotoxic T lymphocytes (CD45^+^CD3^+^CD8^+^ cells; f) and T helper lymphocytes (CD45^+^CD3^+^CD4^+^ cells; g) are unaffected by incorporation of metformin in aPLA or cPLA implants. **h-i**, Unlike with aPLA implants, both T helper 2 cells expressing interleukin-4 (IL-4; h) as well as T helper 1 cells expressing interferon-gamma (IFN-γ; i) are increased by incorporation of metformin in cPLA implants. Mean (SD), one-way ANOVA followed by Tukey’s multiple comparison test, n = 3.

Sterile polyethylene and hydroxyapatite particles, like bacterial lipopolysaccharide (endotoxin), are agonists of Toll-like receptor 4^8,36^. Following activation, by endotoxin or polyethylene particles, increased oxygen consumption at complex I results in the generation of reactive oxygen species that drive proinflammatory activation^20,21^. With respect to endotoxin or polyethylene particles, metformin’s capacity to target complex I results in immunomodulatory effects^21,22,37^. However, unlike endotoxin and polyethylene particles, PLA degradation products signal by binding to monocarboxylate transporters^1,2,38^, which may account for why we did not observe immunomodulatory effects by metformin. In regenerative medicine, such as with musculoskeletal tissues, the inclusion of metformin in biomaterial is beneficial through metformin’s ability to enhance osteoblastogenesis^39-41^. Thus, it is possible that subsequent crosstalk between surrounding immune cells and metformin-activated osteoblasts may occur, necessitating further interrogation. Unlike PLA degradation products that increase cellular ATP levels^1,2^, endotoxin alone^20^ as well as sterile or endotoxin-contaminated polyethylene particles reduce cellular ATP levels^8,21^, suggesting a different metabolic phenotype. While macrophages activated by endotoxin or polyethylene particles rely on glycolysis and not mitochondrial respiration for ATP^8,20^, our findings demonstrate that macrophages exposed to aPLA degradation products rely on mitochondrial respiration for ATP production, suggesting that metabolic rewiring by PLA breakdown products is unique in many respects.

## Materials and Methods

### Polylactide (PLA) materials and extraction

Semi-crystalline PLA 3100HP and amorphous PLA 4060D (both from NatureWorks LLC) were used after confirming their physical, chemical and thermal properties as previously reported^1^. Prior to use, PLA was sterilized by ultraviolet radiation for 30 minutes^42^. Extracts (degradation products)^43^ of PLA were supernatants collected after suspending 4 g of PLA in 25 mL of full medium. Full medium was made by adding 10% heat-inactivated Fetal Bovine Serum and 100 U/mL penicillin-streptomycin to DMEM medium (all from ThermoFisher Scientific). The extraction process lasted 12 days in an orbital shaker (set at 250 rpm and 37 °C). We have previously characterized the monomers and oligomers of lactic acid present in these extracts as well as the pH of extracts^1^. The volume of extract used during in-vitro experiments is noted in respective figure legends.

### Macrophage culture

Mouse primary bone-marrow derived macrophages (BMDMs) were isolated^44^ from male and female 3-4 month old C57BL/6J mice (Jackson Laboratories). Generally, a total of 200 uL of complete medium was used to seed 50,000 BMDMs per well of a 96-well plate for experiments.

### Inhibitors of mitochondrial respiration

Rotenone, metformin, oligomycin and antimycin A were obtained from MilliporeSigma and used at concentrations shown in each figure for probing mitochondrial functions as previously described^21^. Each inhibitor was reconstituted in complete medium and added when seeding cells for in-vitro experiments.

### ATP assessment

We used ATP/ADP kits (Sigma-Aldrich) containing D-luciferin, luciferase and cell lysis buffer according to manufacturer’s instructions to assess ATP levels in cultured BMDMs. Relative Luminescence Units (RLU) measurements were collected at integration time of 1000 ms using the SpectraMax M3 Spectrophotometer (Molecular Devices) and SoftMax Pro software (Version 7.0.2, Molecular Devices).

### Macrophage viability

We assessed BMDMs viability using the crystal violet assay^45^, where absorbance (optical density) was acquired at 570 nm using the the SpectraMax M3 Spectrophotometer (Molecular Devices) and SoftMax Pro software (Version 7.0.2, Molecular Devices).

### Seahorse assay on cultured macrophages

Basal measurements of oxygen consumption rate (OCR) were acquired with the XFe-96 Extracellular Flux Analyzer (Agilent Technologies). We used the Seahorse ATP rate assay according to manufacturer’s instructions. Following cell culture in complete medium, before running the Seahorse assay, we washed and cultured BMDMs in the Seahorse XF DMEM medium (pH 7.4) supplemented with 25 mM D-glucose and 4 mM Glutamine (all sourced from Agilent Technologies) for an hour in a non-CO2 incubator at 37 °C. To export data as mean (SD), Wave software (Version 2.6.1) was used.

### Mitochondrial reactive oxygen species (ROS) measurement

As previously reported^21^, cultured primary bone marrow-derived macrophages were separated from 96-well plates using 4 mM EDTA (Teknova) in 1X PBS at 37 °C for 10 min, combined with gentle pipetting. Macrophages were evaluated for mitochondrial mass (MitoTracker Green FM, MTG; ThermoFisher Scientific) and mitochondrial superoxide (MitoSOX Red, MSOX; ThermoFisher Scientific). We used unstained cells from each condition and single-stained MTG and MSOX cells as controls for gating. A cocktail of 1X PBS (ThermoFisher Scientific) containing Zombie NIR (1:750), MTG (50 nM) and MSOX (2.5 μM) was added to samples with 40,000 cells in 100 μL, followed by incubation at 37 °C and 5% CO2 in the dark for 30 min. Afterwards, cells were collected by centrifugation followed by two washes using flow buffer (0.5% bovine serum albumin (MilliporeSigma) in 1X PBS), then fixed using 4% paraformaldehyde for 10 min at room temperature in the dark. Fixed cells were collected by centrifugation then suspended in 100 μL flow buffer for analysis using the Cytek Aurora flow cytometer. Cells were identified and singlets gated using FSC and SSC. While MTG^+^ cells were gated from live cells, MSOX^+^ cells were identified from the MTG^+^ population. All flow cytometry data were analyzed using FCS Express software (De Novo Software; version 7.12.0035).

### Cytokine and Chemokine measurements

As previously reported^1,2,8,21^, cytokine and chemokine levels in cell culture supernatants were measured by a MILLIPLEX MAP mouse magnetic bead multiplex kit (MilliporeSigma) specifically for IL-6, MCP-1, TNF-α, IL-1β, IL-4, IL-10, IFN-γ and 1L-13. We obtained the data using a Luminex 200 instrument (Luminex Corporation) and xPONENT software (Version 3.1, Luminex Corporation).

### Fabrication of biomaterials incorporating inhibitors

Metformin (MilliporeSigma) was incorporated into amorphous PLA (aPLA) or crystalline PLA (cPLA) by melt-blending at 190 ºC for 3 mins in a DSM 15 cc mini-extruder (DSM Xplore), then made into pellets using a pelletizer (Leistritz Extrusion Technology) as previously described^1,5^. Thereafter, 1.75 mm (diameter) filaments were extruded using a Filabot EX2 at 170 ºC with air set at 93. To estimate 10 mM metformin applied in-vitro, we added 12.916 mg of metformin to 9.987g of aPLA or cPLA. Extruded filaments were cut into 1 mm-long sizes, then sterilized by ultraviolet radiation for 30 mins before surgical implantation. Both aPLA and cPLA were reprocessed under similar conditions, without the addition of metformin, to control for melt-blending as a confounder in our studies. This method has been previously validated to load small molecules, whose release profiles have been characterized, onto aPLA and cPLA implants^5,10^.

### Animal studies

Experiments involving subcutaneous biomaterial implantation in mice were approved by the Institutional Animal Care and Use Committee at Michigan State University (PROTO202100327). Female C57BL/6J (wild-type B6) mice (9-week old) were obtained from the Jackson Laboratory, with n = 3 mice assigned to each study group. Under anaesthesia, by isoflurane (2-3 %), the dorsum of mice was shaved and disinfected using alternate iodine and alcohol swabs. Afterwards, incisions were made through the skin into the subcutis, without (sham groups) or with biomaterial implantation as previously described^5^. We used surgical glue (3M Vetbond) to close the incision, and mice received postoperative analgesia by meloxicam (5 mg/ kg) administered subcutaneously.

### Tissue dissociation for flow cytometry

Mice were shaved around the incision areas (sham) or biomaterials, then euthanized to collect circular biopsies (8 mm diameter). We pooled tissues from different mice belonging to the same group for dissociation using a combination of mechanical and enzymatic dissociation strategies. Tissues were dissected into small sections and rubbed on a serrated Petri dish containing 10 mL of an enzyme cocktail comprising 0.5 mg/ mL Collagenase Type IV (Stem Cell Technologies), 0.5 mg/ mL Liberase (Sigma-Aldrich) and 250 U/ mL Deoxyribonuclease I (Worthington Biochemical Corporation) in 25mM HEPES buffer (Sigma-Aldrich). Enzymatic digestion was undertaken on an orbital shaker (70 rpm) within a 5% CO2 incubator at 37 °C for an hour. Afterwards, the 10 mL enzyme cocktail was filtered through a 70 μm filter into a 50mL conical tube. Undigested tissues were further pressed against serrated Petri dishes or with the thumb press of a syringe plunger. We used cold Hanks’ Balanced Salt Solution without calcium, magnesium and phenol red (ThermoFisher Scientific) to wash dissociated cells off Petri dishes, followed by another cycle of filtration through a 70 μm filter. Centrifugation at 350 x g for 10 minutes allowed for sedimentation of cells, which were counted for flow cytometry.

Following tissue digestion, dissociated cells were split into n = 3, each containing 1×10^6^ cells for staining in a polypropylene 96-well round bottom plate. Staining steps were performed in 100 μL volume in the dark at 4DC. We incubated samples with LIVE/DEAD Fixable Blue Dead Cell Stain Kit (1:500, Thermofisher, cat#L23105) for 20 minutes. Thereafter, cells were washed once with flow buffer and incubated with TruStain FcX (anti-mouse CD16/32) Antibody (BioLegend, 101319; 1 μg/sample) in 50 μL volume for 10 minutes. We mixed the following antibodies and added them to the cell suspension for a 30 mins incubation: BV605 CD45 (1:500, Biolegend, 103139), BV785 F4/80 (1:300, Biolegend, 123141), BV421 CD86 (1:200, Biolegend, 105031), APC CD206 (1:200, Biolegend, 141707), PerCP MHCII (1:200, Biolegend, 107623), SparkBlue 550 CD3 (1:100, Biolegend, 100259), APC-Fire 810 CD4 (1:100, Biolegend, 100479), BB700 CD8a (1:100, Biolegend, 566410), and PE-Dazzle 594 CD11c (1:500, Biolegend, 117347). Before fixation and permeabilization (BD Cytofix/Cytoperm kit, BDB554714), cells were washed once, suspended in BD Perm/wash buffer with BV650 IL4 (1:50, BD Bioscience, cat#564004), APC-Fire750 IFNγ (1:80, Biolegend, 505859), and PE-Cy7 Arg1 (1:100, ThermoFisher, 25-3697-80) for 30 mins incubation. Thereafter, cells were washed twice with BD Perm/wash buffer then suspended in a final volume of 100 μL for flow cytometry analysis. The Cytek Aurora spectral flow cytometer (Cytek Biosciences, CA, USA) was used for sample analyses, using the Cytek SpectroFlo software (version 3.0.3) for data collection. Fluorescence minus one (FMO) samples were used to guide gating strategies, and the flow cytometry data was analyzed with the software FCSExpress (DeNovo Software, CA, USA; version 7.12.0005).

### Statistics

Statistics was undertaken using software (GraphPad Prism Version 9.5.1 (528)). Analyzed data were presented as mean with standard deviation (SD), and the exact statistical test, p-values and sample sizes are provided in figure legends.

## Data availability

The data regarding the findings of this study are available within the paper.

## Acknowledgements

Funding for this work was provided in part by the James and Kathleen Cornelius Endowment at MSU.

## Author contributions

Conceptualization, C.V.M. and C.H.C.; Methodology, C.V.M., A.V.M., A.T., E.U., K.B.S., M.A., R.N., S.B.G., N.A., J.H.E., K.D.H. and C.H.C.; Investigation, C.V.M., A.T., A.V.M., E.U.; Writing – Original Draft, C.V.M.; Writing – Review & Editing, C.V.M., A.V.M., A.T., E.U., K.B.S., M.A., R.N., S.B.G., N.A., J.H.E., K.D.H. and C.H.C.; Funding Acquisition, C.H.C.; Resources, R.N. and C.H.C.; Supervision, R.N., J.H.E., K.D.H. and C.H.C.

## Competing interests

The authors declare no competing interest.

## Notes

### Competing Interest Statement

The authors have declared no competing interest.

